# Acute restraint stress and pain modulation depend on the interaction between the periaqueductal gray and the lateral septum

**DOI:** 10.1101/2024.11.29.626041

**Authors:** Devanshi Piyush Shah, Yatika Chaudhury, Arnab Barik

## Abstract

Acute restraint stress is known to cause analgesia in humans and laboratory animals, but the mechanisms are unknown. Recently, we have shown that a multi-nodal circuitry between the dorsal lateral septum (dLS)-lateral hypothalamic area (LHA)-rostral ventromedial medulla (RVM) plays an instructive role in restraint stress-induced analgesia. We found that the LS neurons are activated when mice struggle to escape the restraint, and we wondered about the origin of the escape signals. Hence, we performed retrograde viral labeling from the LS and found that the ventrolateral periaqueductal gray (vlPAG), a known anatomical substrate for escape behaviors, provides inputs to the LS. Through anatomical, behavioral, and in-vivo fiber photometry, we show that the PAG and LS neurons are synaptically connected; activation of either PAG or the post-synaptic LS neurons is sufficient to cause analgesia and sufficiently cause hyperalgesia. Moreover, we found that the LS neurons that receive inputs from PAG send axonal projections to the LHA. Together, we found that the vlPAG neurons encoding nociceptive and escape behaviors provide synaptic inputs to the dLS-LHA-RVM circuitry to mediate acute restraint stress-induced analgesia.

**Significance statement:** Forced restraint causes stress, and this paradigm has been used in the laboratory to study the physiological effects of stress, including pain modulation. The dorsal lateral septum (LS) has been shown to mediate restraint-mediated stress and analgesia. It is unknown what signals encoded in the LS allow it to instruct stress and pain. A novel neural pathway between the periaqueductal gray (PAG) and LS is activated when mice try to escape the restraint. Repeated activity in the PAG-LS circuitry due to the inescapability of the restraint causes stress and analgesia

## Introduction

Stress-induced analgesia (SIA) is a phenomenon where the presence of an organism in an acutely stressful environment leads to pain alleviation (Amit and Galina, 1986; Butler and Finn, 2009; Yilmaz et al., 2010). This mechanism is crucial as it allows organisms to tend to their physiological needs, which are critical for survival. The neural mechanisms that translate acute stress into pain modulation are poorly understood.

Restraint stress (RS) is a commonly used stress-inducing paradigm in laboratory animals. Acute RS activates lateral septum (LS) neurons and causes elevated blood corticosterone levels and anxiety (Anthony et al., 2014; Besnard et al., 2019; Azevedo et al., 2020; An et al., 2022; Shah et al., 2024). The LS neurons mediate acute RS-induced SIA through their inputs to the lateral hypothalamus (LH) (Shah et al., 2024). Notably, the LS neurons were found to be precisely tuned to the escape-like struggles that mice exhibited under restraint—indicating that the inability to escape may precipitate stress and anxiety seen in animals that have undergone RS (Azevedo et al., 2020; Shah et al., 2024). Here, we investigated the circuit mechanisms that inform the LS of the efforts to escape restraint.

The periaqueductal gray (PAG) is a crucial brain nuclei involved in escape behavior and defensive responses (Branco and Redgrave, 2020; Lefler et al., 2020). Seminal electrical and chemical studies dissected the functional topography of the PAG: the stimulation of the dorsomedial and dorsolateral parts evoked jumping and running(Anon, 1988; Lefler et al., 2020). In contrast, the ventrolateral part (vlPAG) caused freezing and immobility (Anon, 1988; Caroline Blanchard et al., 2009; Depaulis and Bandler, 2012). More recent studies have shed light on how the dorsal PAG computes escape behaviors in natural settings (Evans et al., 2018; Reis et al., 2021a, 2021b; Stempel et al., 2024). These findings highlight the PAG’s importance in integrating brain-wide information to generate flexible defensive and motivated behaviors. We found that the PAG neurons send long-distance axonal projections to the LS. The LS neurons downstream of PAG (LS_post-PAG_) are tuned to the bouts of struggle in mice under RS, and the LS_post-PAG_ neurons bidirectionally modulate nociceptive thresholds. Thus, the PAG-LS may be vital in translating inescapable restraint to acute stress to subsequent pain modulation.

## Materials and Methods

### Animals

Animal care and experimental procedures were performed following protocols approved by the CPSCEA at the Indian Institute of Science. The animals were housed at the IISc Central Animal Facility under standard animal housing conditions: 12 h light/dark cycle from 7:00 am to 7:00 pm with ad libitum access to food and water; mice were housed in IVC cages in Specific pathogen-free (SPF) clean air rooms. Mice strains used: *Vglut2^Cre^* or *Vglut2-ires-Cre* or *Slc17a6tm2(Cre)* Lowl/J(Stock number 016963); BALB/cJ (Jackson Laboratories, USA). Experimental animals were between 2-4 months old.

### Viral vectors and stereotaxic injections

Mice were anesthetized with 2% isoflurane/oxygen before and during the surgery. Craniotomy was performed at the marked point using a hand-held micro-drill (RWD, China). A Hamilton syringe (10 μL) with a glass pulled needle infused 300 nL of viral particles (1:1 in saline) at 100 nL/minute. The following coordinates were used to introduce virus/dyes: PAG-Anterior-Posterior (AP): -4.84, Medial-Lateral (ML): ±0.50, Dorsal-Ventral (DV): -2.75; dLS-AP: +0.50, ML: + 0.25; DV: -2.50; and LHA-AP: -1.70, ML: ±1.00; DV: -5.15. Vectors used and sources: AAV9.syn.flex.GcaMP6s (Addgene, Catalog# pNM V3872TI-R(7.5)), AAVretro-hSyn-NLS-mCherry (Donated by Ariel Levine, NIH), AAV5-hSyn-DIO-mSyp1_EGFP (University of Zurich, Catalog# v484-9), AAV9-DIO-PSD95-TagRFP (Donated by Mark Hoon, NIH), AAV1-hSyn-Cre.WPRE.hGH (Addgene, Catalog# v126225), pAAV5-hsyn-DIO-hM3D(Gq)-mCherry (Addgene, Catalog# v141469), pAAV-Ef1a-DIO-eNPHR 3.0-EYFP (Addgene, Catalog# v32533), pAAV5-hsyn-DIO-EGFP (Addgene, Catalog# 50457-AAV 1), pENN.AAV5.hSyn.TurboRFP.WPRE.RBG (Addgene, Catalog# 10552-AAV1).

For the rabies tracing experiments, rAAV5-EF1α-DIO-oRVG (BrainVTA, Catalog# PT-0023) and rAAV5-EF1α-DIO-EGFP-T2A-TVA (BrainVTA, Catalog# PT-0062) were injected first, followed by RV-EnvA-Delta G-dsRed (BrainVTA, Catalog# R01002) after two weeks. Tissue was harvested after one week of rabies injection for histochemical analysis. Post-hoc histological examination of each injected mouse was used to confirm that viral-mediated expression was restricted to target nuclei.

### Fiber implantation

Fiber optic cannula from RWD, China; Ø1.25 mm Ceramic Ferrule, 200 μm Core, 0.22 NA, L = 5 mm were implanted at AP: -4.84, ML: +0.50; DV: -2.75 in the vlPAG; and at AP: 0.50, ML: +0.25; DV: -2.50 in the dLS after AAV carrying GCaMP6s or Halorhodopsin were infused. Animals were allowed to recover for at least three weeks before performing behavioral tests. Before every assay, the fibers were connected to the cannulae, and the mice were allowed to acclimatize to them for 30 minutes. Successful labeling and fiber implantation were confirmed post hoc by staining for GFP for viral expression and injury caused by the fiber for implantation. Animals with viral-mediated gene expression at the intended locations and fiber implantations, as observed in post hoc tests, were only included.

### Behavioral assays

A single experimenter handled behavioral assays for the same cohorts. Before experiments, mice were habituated in their home cages for at least 30 minutes in the behavior room. An equal male-to-female ratio was maintained in every experimental cohort and condition unless otherwise stated, with no significant differences seen between sexes in the responses recorded from the behavioral experiments. Wherever possible, efforts were made to keep the usage of animals to a minimum.

The mice were subjected to five behavioral assays. In the immobilization experiments, experimenters physically restrained the mice by pressing them down by hand for approximately 10 seconds. In the tail-hanging experiments, the mice were suspended upside down by their tail for 10 seconds. In the restraint stress (RS) assay, photometry signals were recorded through the fiber-coupled cannulae that passed through a modified RS-inducing falcon tube to allow unrestricted recording (Shah et al., 2024). On the gradient hot plate test (Orchid Scientific, India), the mice were subjected to a range of temperature from 32-52 degrees for 5 minutes, with the temperature increasing at a rate of 4.8 degrees/minute on a hot plate surrounded by an inescapable transparent acrylic enclosure to enable video monitoring (Logitech, USA). In the tail-flick assay (Orchid Scientific, India), an intense light beam is focused on the mice’s tails until they flick it away.

These assays were performed for chemogenetic stimulation of the neurons, optogenetic inhibition of the neurons, and fiber photometry to record the neural transients. Chemogenetic stimulation of neurons was brought about by expressing DREADD-carrying viruses in specific neurons of interest and an i.p injection of the ligand Deschloroclozapine (DCZ) (diluted in saline to a final concentration of 0.1 mg/kg) 15-20 minutes before behavioral experiments or histochemical analysis. Optogenetic inhibition of neurons was brought about by expressing inhibitory opsin (halorhodopsin)-carrying viruses in the neurons of interest and shining yellow light through fiber cannulae implanted in the desired co-ordinates of the brain (Prizmatix, Israel). A dual-channel fiber photometry system from RWD (R810 model) was used to record the data for fiber photometry experiments. The light from two light LEDs (410 and 470 nm) was passed through a fiber optic cable coupled to the cannula implanted in the mouse. Fluorescence emission was acquired through the same fiber optic cable onto a CMOS camera through a dichroic filter. The photometry data was analyzed using the RWD software, and .csv files were generated. The start and end of stimuli were timestamped. All trace graphs were plotted from .csv files using GraphPad Prism software version 8.

### Immunostaining, multiplex in situ hybridization, and confocal microscopy

Mice were anesthetized with isoflurane and perfused intracardially with 1X Phosphate Buffered Saline (PBS) (Takara, Japan) and 4% Paraformaldehyde (PFA) (Ted Pella, Inc., USA), consecutively for immunostaining experiments. Tissue sections were rinsed in 1X PBS and incubated in a blocking buffer (2% Bovine Serum Albumin (BSA); 0.3% Triton X-100; PBS) for 1 hour at room temperature. Sections were incubated in primary antibodies made in the blocking buffer at room temperature overnight. Sections were rinsed 1-2 times with 1X PBS and incubated for 2 hours in Alexa Fluor conjugated goat anti-rabbit/ chicken or donkey anti-goat/rabbit secondary antibodies (Invitrogen) at room temperature, washed in 1X PBS, and mounted in VectaMount permanent mounting media (Vector Laboratories Inc.) onto charged glass slides (Globe Scientific Inc.). We used an upright fluorescence microscope (Khush, Bengaluru) (2.5X, 4X, and 10X lenses) and ImageJ/FIJI image processing software to image and process images to verify the anatomical location of cannulae implants. For the anatomical studies, the images were collected with 10X and 20X objectives on a laser scanning confocal system (Leica SP8 Falcon, Germany) and processed using the Leica image analysis suite.

### Quantification and Statistical Analysis

All statistical analyses (t-test and one-way ANOVA test) were performed using GraphPad PRISM 8.0.2 software. ns > 0.05, ∗ *P* ≤ 0.05, ∗∗ *P* ≤ 0.01, ∗∗∗ *P* ≤ 0.001, ∗∗∗∗ *P* ≤ 0.0005.

### Data availability

Raw data will be made available upon request.

## Results

### Projection neurons from vlPAG neurons form excitatory synapses on dLS neurons

The dorsal part of the LS encodes bouts of struggle or efforts to escape. Under acute restraint, the inability to escape results in stress and analgesia (Shah et al., 2024). Acute activation of LS causes stress and anxiety however, do not elicit escape behaviors. Hence, the efforts to escape under acute restraint are likely encoded in the pre-synaptic partners of the LS. Here, we sought to understand the synaptic inputs of LS neurons that may inform them of the struggles or efforts to escape in the acute RS assay. To this end, we injected the retrograde viral tracer AAVRetro-NLS-mCherry into the dLS to map the upstream brain regions projecting onto the inhibitory neurons of the dLS (Figures 1A, B). The mCherry, fused to a nuclear localization signal (NLS), ensures nuclear-specific expression of the fluorescent protein. During anatomical visualization of neurons, the nuclear localization of fluorescent protein allows clear identification of the retrograde pre-synaptic targets (Sathyamurthy et al., 2020; Anon, 2022). Interestingly, we found neurons labeled with mCherry in the ventrolateral region of the periaqueductal gray (vlPAG) (Figure 1C). The PAG is comprised of both excitatory and inhibitory neurons. We hypothesized that the glutamatergic neurons project to the dLS, suggesting that the escape responses encoded in the PAG can consistently engage the dLS neurons, resulting in SIA. Thus, we labeled the glutamatergic neurons in the vlPAG with synaptophysin-tagged GFP (SypGFP) to enable visualization of the synaptic vesicle-rich areas in the axon terminals at target sites such as dLS. To that end, we stereotaxically injected the Cre-dependent DIO-SypGFP virus in the vlPAG of the VGlut2-Cre (vlPAG^VGlut2^) transgenic mice (Figure 1D). We observed Syp-GFP puncta in the dLS, indicating that the vlPAG^VGlut2^ neurons synapse with the inhibitory neurons in the dLS (Figure 1E). To confirm if the vlPAG^VGlut2^ neurons form synapses with the inhibitory dLS neurons, we expressed the excitatory post-synaptic density protein, PSD-95, tagged with a fluorescent protein, tagRFP (PSD95tagRFP) driven by the *Gad1* promoter, in the inhibitory dLS (dLS*^Gad1^*) neurons (Figure 1D). Thus, in the same VGlut2-Cre mice that were injected with AAV-DIO-SypGFP in the vlPAG, we injected a mixture of AAV-*Gad1*-Cre and DIO-PSD95tagRFP in the dLS (Figure 1D)—allowing simultaneous labeling the pre-synaptic and post-synaptic structures with fluorescent proteins in non-overlapping spectra. We found that the synaptic terminals of the excitatory projections from the vlPAG^VGlut2^ neurons (green) were closely apposed with the post-synaptic compartments of the dLS*^Gad1^ cells* (red) (Figure 1E). This suggests that excitatory neurons from the vlPAG synapse onto inhibitory neurons in the dLS.

**Figure 1.**
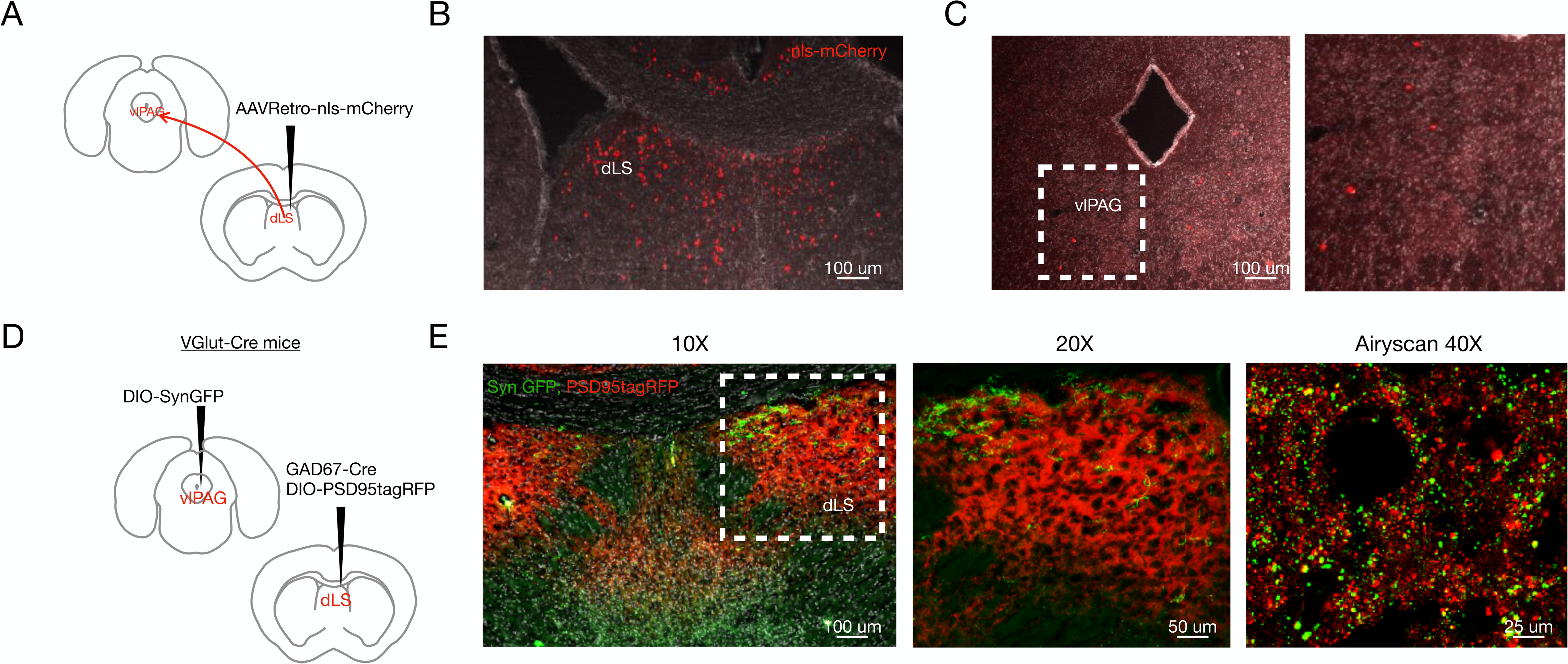
Excitatory vlPAG neurons synapse onto inhibitory dLS neurons. (A) RetroAAV-nlsmCherry was injected in the dLS of wild-type mice to trace cell bodies of presynaptic neurons in the vlPAG. (B) Confirmation of the expression of nls-mCherry (red) in the dLS. (C) Red cell bodies were seen in the vlPAG (marked region zoomed-in on the right). (D) DIO-SynGFP was injected in the vlPAG, and DIO-PSD95-tagRFP in the dLS of VGlut2-Cre transgenic mice. (E) Confocal images of closely apposed green (vlPAG^VGlut2^ terminals) and red (dLS^VGlut2^ dendrites) cells were observed under the confocal microscope at 10X and 20X magnifications and 40X under the Airyscan.

### vlPAG^VGlut2^ neurons respond to escape behaviors during the RS assay

Since the dLS*^Gad1^* neurons have been known to be engaged during acute-restraint stress (Shah et al., 2024), and vlPAG^VGlut2^-dLS*^Gad1^* neurons are synaptically connected (Figure 1E), we sought to test whether the vlPAG^VGlut2^ neurons are activated when the mice are under acute restraint-stress. We expressed GCaMP6s in the vlPAG^VGlut2^ neurons by delivering AAV-DIO-GCaMP6s in the vlPAG of VGlut2-Cre transgenic mice (Figures 2A, B). We recorded the calcium transients from the GCaMP6s-labeled neurons using fiber photometry while the mice were subjected to RS (Figure 2C). Neural recordings during the RS assay showed that vlPAG^VGlut2^ neurons were engaged when the mice were acutely restrained (Figure 2D), with peaks in activity explicitly aligning with the bouts of the mice struggling to escape in the experimental tube (Figure 2E). Next, we tested if the vlPAG^VGlut2^ neurons respond to noxious thermal stimuli. We used the gradient thermal plate assay to expose the mice to increasing levels of surface temperatures (Figure 2F) (Espejo and Mir, 1993; Barik et al., 2018; Reddy et al., 2023). In this assay, mice are placed on the thermal plate, surrounded by an inescapable transparent acrylic enclosure to enable video monitoring (Figure 2F). When the temperature of the thermal plate crosses the nociceptive thresholds (above 44°C), the mice tend to shake and lick their paws to cope with the heat pain. When the temperature crosses 52°C, the mice jump to escape (Figures 2G, H). The vlPAG^VGlut2^ neurons started responding once the hot-plate temperature crossed 44°C, and the peaks in the activity corresponded with the number of jumps (Figures 2G, H). Implying that noxious heat engages vlPAG^VGlut2^ neurons, suggesting that these cells are tuned to escape responses on the hot-plate test.

**Figure 2.**
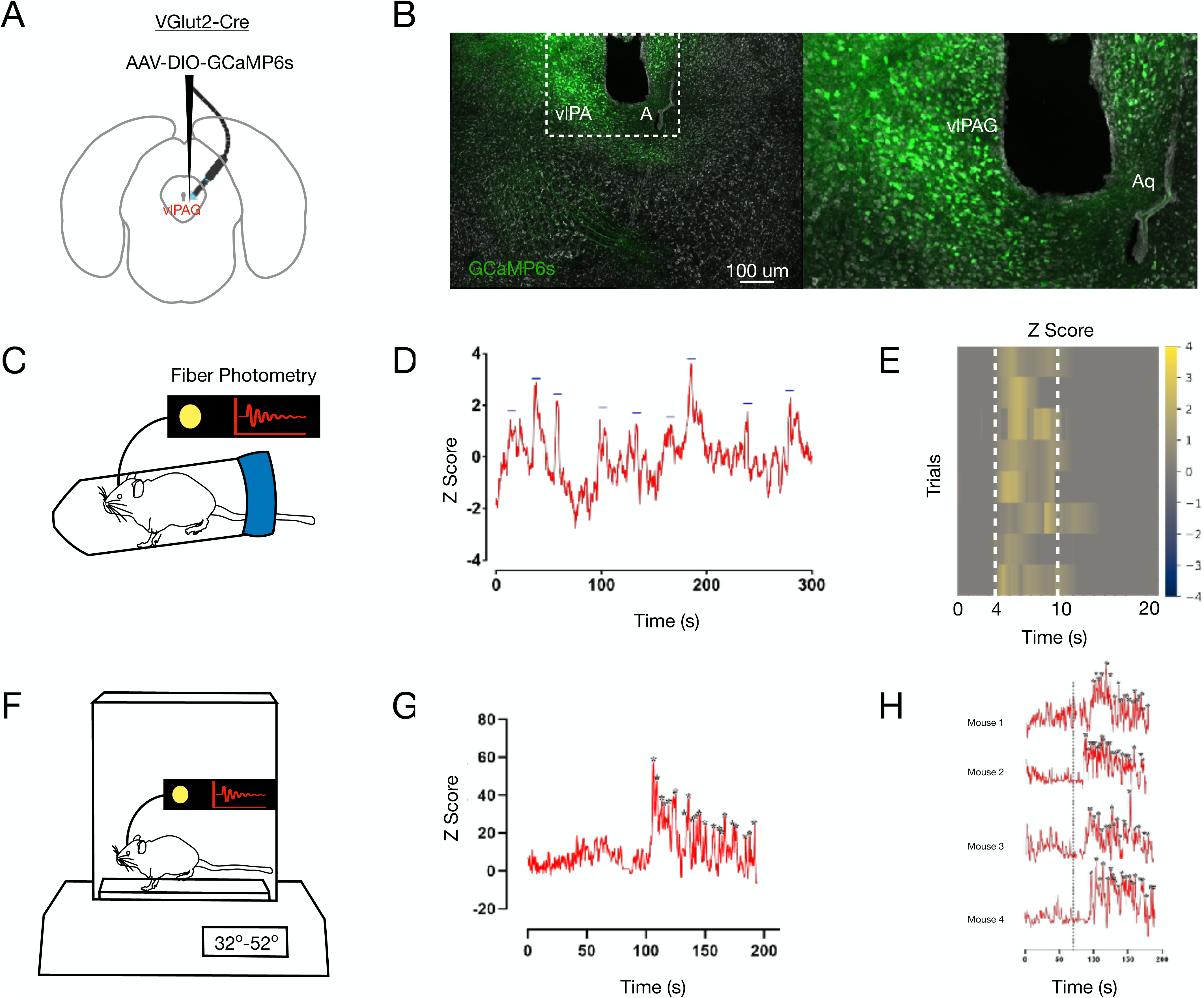
vlPAG^VGlut2^ neurons respond to struggle during RS assay. (A) AAV encoding DIO-GCaMP6s was injected in the vlPAG of VGlut2-Cre transgenic mice, and fiber optic cannulae were implanted at the same coordinates for recording neural activity. (B) Confirmation of GCaMP6s (green) expression in the vlPAG neurons (marked by white box). (C) Schematic representation of fiber photometry recording during the RS assay. (D) Representative trace of neural activity from vlPAG^GCaMP6s^ neurons from mice under restraint in the RS assay (blue dashes indicate neural activity during struggle behavior in the RS assay). (E) Heatmap depicting neural activity during individual struggles (initiation and end of struggle indicated by white dotted lines, n=5). (F) Schematic representation of fiber photometry recording during the hot plate assay. (G) Representative trace of the neural activity from vlPAG^GCaMP6s^ neurons from mice on the gradient hotplate at 32-52 degrees (stars indicate neural activity corresponding to jumping behavior on the hotplate). (H) Traces of neural activity from vlPAG^GCaMP6s^ neurons from all mice on the gradient hotplate at 32-52 degrees (stars indicate neural activity corresponding to jumping behavior on the hotplate, initiation of noxious temperature indicated by black dotted line) (n=5).

### dLS_post-vlPAG_ neurons are activated during the struggle in the RS assay

We found that the vlPAG^VGlut2^ neurons are tuned to anxiogenic and high-threshold thermal stimuli that evoke escape behaviors (Figures 2G, H). Next, we sought to understand how the downstream neurons in the LS (dLS_post-vlPAG_) respond to restraint stress and noxious thermal stimuli. Before this, dLS neuronal activity correlated with stressors such as tail hanging, immobilization, and acute restraint (Azevedo et al., 2020; Shah et al., 2024). In the RS assay, the bouts of struggle to escape correlated with the dLS activity (Shah et al., 2024). Hence, we labeled the dLS_post-vlPAG_ neurons with GCaMP6s by injecting the anterograde transsynaptic AAV1-hSyn-Cre (AAVTranssyn-Cre) in the vlPAG and the Cre-dependent GCaMP6s in the dLS (Figures 3A, B). Then, we recorded the activity from the dLS_post-vlPAG_ neurons with fiber photometry as the mice were exposed to acute stressors and noxious thermal stimuli. dLS_post-vlPAG_ neurons responded to the struggle bouts in the RS assay, as observed in the pre-synaptic vlPAG neurons and previous recordings from the dLS (Figures 3C, D). On the gradient hot-plate assay, the dLS_post-vlPAG_ neurons responded once the plate temperature crossed the nociceptive thresholds into the unbearable range (Figures 3E, F). Similarly, hand immobilization and tail-hanging, two commonly used stressors, activated the dLS_post-vlPAG_ neurons (Figures 3G-J). This finding contrasts with the recordings from the dLS neurons, where in fiber photometry recordings, noxious heat failed to elicit a response from the dLS neurons. This discrepancy might be due to recording from a more specific neural population here. A small proportion of neurons, including the dLS_post-vlPAG_ neurons, are tuned to noxious thermal stimuli (Figures 3E, F). Moreover, as seen in the pre-synaptic vlPAG^VGlut2^ neurons, the rise in activity of the dLS_post-vlPAG_ neurons correlated with escape bouts on the gradient hot-plate test (Figures 2G, H). Thus, the dLS_post-vlPAG_ neurons are tuned to anxiogenic stressors and noxious thermal behavior with the potential to stimulate escape behaviors.

**Figure 3.**
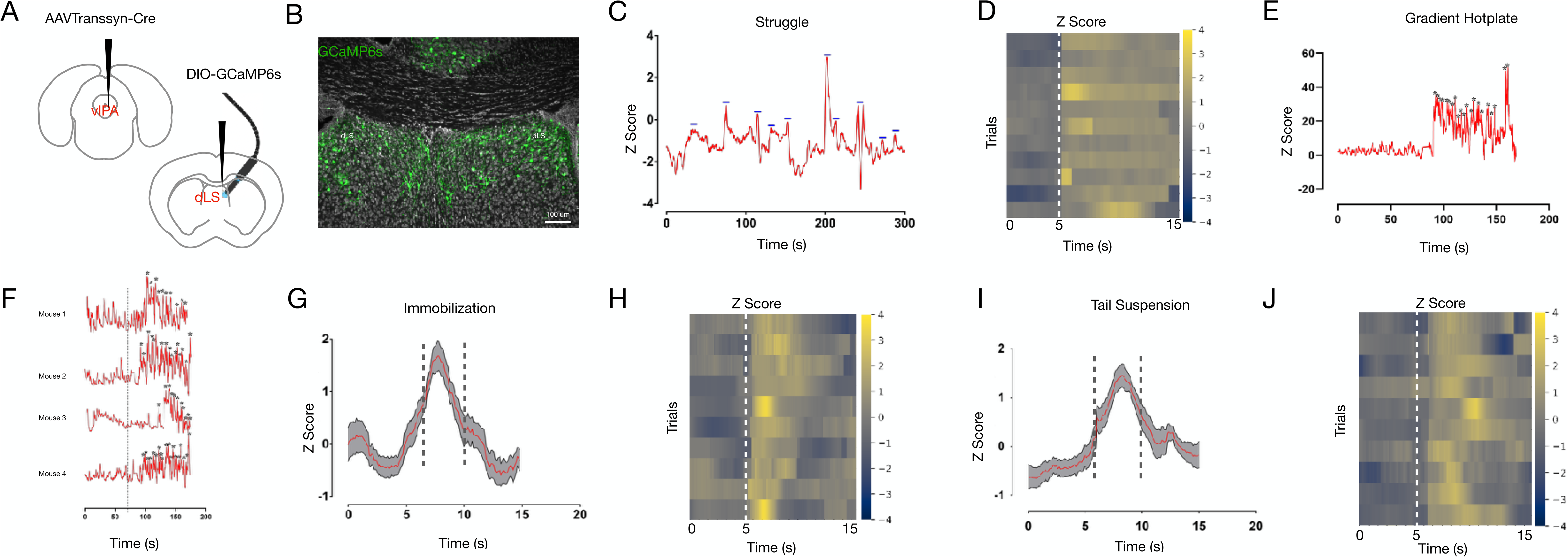
dLS_post-vlPAG_ neurons are activated during the struggle in the RS assay. (A) AAVTranssyn-Cre was injected in the vlPAG and DIO-GCaMP6s in the dLS of wild-type mice, and fiber optic cannulae were implanted at the dLS for recording neural activity. (B) Confirmation of the expression of GCaMP6s (green) in the dLS neurons. (C) Representative trace of neural activity from dLS^post-vlPAG^ neurons in mice under restraint in the RS assay (blue dashes indicate neural activity during struggle behavior in the RS assay). (D) Heatmap depicting neural activity during individual struggles (initiation of struggle indicated by a white dotted line, n=5). (E) Representative traces of neural activity from dLS^post-vlPAG^ neurons from mice on the gradient hotplate at 32-52 degrees (stars indicate neural activity corresponding to jumping behavior on the hotplate). (F) Traces of neural activity from dLS^post-vlPAG^ neurons from all mice on the gradient hotplate at 32-52 degrees (stars indicate neural activity corresponding to jumping behavior on the hotplate and initiation of noxious temperature indicated by black dotted line) (n=5). (G and H) Average plot and heatmap (individual trials) of Z-Score trace from dLS^post-vlPAG^ neurons during immobilization (n=5, 16 trials). (I and J) Average plot and heatmap (individual trials) of Z-Score trace from dLS^post-vlPAG^ neurons during tail suspension (n=5, 18 trials).

### dLS_post-vlPAG_ neurons bi-directionally modulate thermal nociceptive thresholds

Next, we transiently activated the dLS_post-vlPAG_ neurons and tested their roles in RS-induced SIA. For the chemogenetic neuronal stimulation, we expressed the activating DREADD actuator, hM3Dq (Roth, 2016), in the dLS_post-vlPAG_ neurons (Figures 4A, B). The hM3Dq neurons are activated by the intra-peritoneal (i.p.) administration of the potent hM3Dq ligand deschloroclozapine (DCZ) (Nagai et al., 2020). We injected AAVTranssyn-Cre in the vlPAG and the Cre-dependent DIO-hM3Dq-mCherry in the LS (Figure 4A). Expression of hM3Dq was confirmed by the mCherry expression in the dLS_post-vlPAG_ neurons (Figure 4B). dLS_post-vlPAG_ stimulation by i.p. DCZ reduced the number and increased the latency to lick on the hot-plate test, set at 52°C (Figure 4C). On the tail-flick test, stimulation of the dLS_post-vlPAG_ neurons increased the latency to react (Figure 4D). Thus, the chemogenetic activation of the dLS_post-vlPAG_ neurons was sufficient to result in the suppression of nociceptive thresholds or analgesia in response to noxious thermal stimuli in the tail-flick test involving spinal-reflex circuitry, as well as the hot-plate test that requires supraspinal participation.

**Figure 4.**
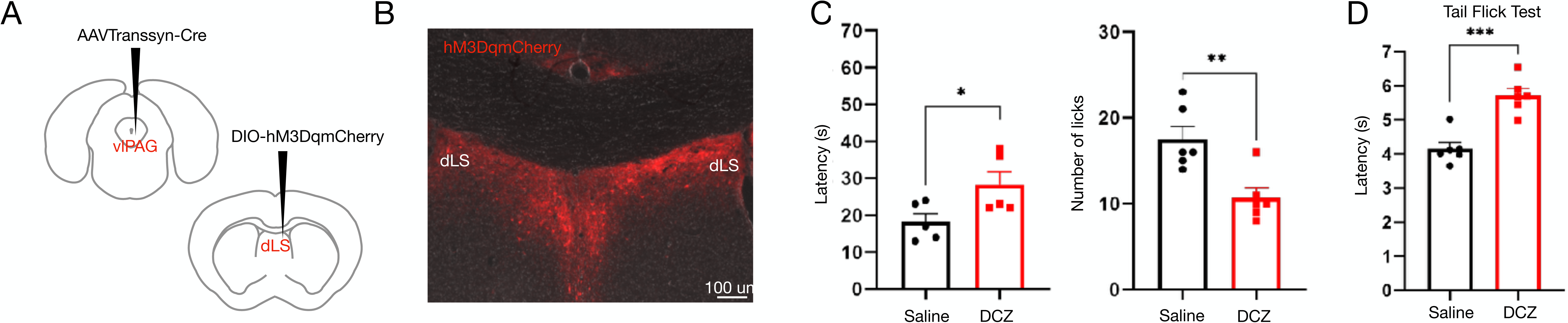
Activation of dLS_post-vlPAG_ neurons is analgesic. (A) AAVTranssyn-Cre was injected in the vlPAG and DIO-hM3Dq-mCherry in the dLS of wild-type mice. (B) The expression of hM3Dq-mCherry (red) was confirmed in the dLS^post-vlPAG^ neurons. (C) Latency to lick (seconds) (17.83±1.88 compared to 27.67±2.99, respectively; t-test, \**P*=0.019, n=6) and licks (17.50±1.47 compared to 10.67±1.14, respectively; t-test, \*\**P*=0.0044, n=6) on the hot-plate test post i.p. Saline or DCZ administration. (D) Tail-flick latency (seconds) (4.16±0.19 compared to 5.71±0.21, respectively; t-test, \*\*\**P*=0.0003, n=6) post-saline or DCZ administration.

The inhibitory light-sensitive channel, Halorhodopsin (eNpHR3.0) (Zhang et al., 2007; Tye et al., 2011), when expressed in the neural population of choice and, upon shining yellow light, was found to hyperpolarize neurons. Thus, we tested if the transsynaptic expression of the eNpHR3.0 in the dLS_post-vlPAG_ neurons would alter the thermal nociceptive thresholds. We stereotaxically injected the AAVTrannsyn-Cre in the vLPAG and the DIO-eNpHR3.0-YFP in the LS (Figure 5A). We confirmed the eNpHR3.0 expression in the dLS_post-vlPAG_ neurons by visualizing the YFP (Figure 5B). We found that by inactivating the dLS_post-vlPAG_ neurons decreased nociceptive thresholds to paw-licking and tail-flick behaviors and increased nocifensive licking behavior to noxious thermal stimuli (Figures 5C, D). Therefore, dLS_post-vlPAG_ neurons can bi-directionally regulate thermal nociceptive thresholds and noxious heat-induced spinal and supraspinal defensive behaviors.

**Figure 5.**
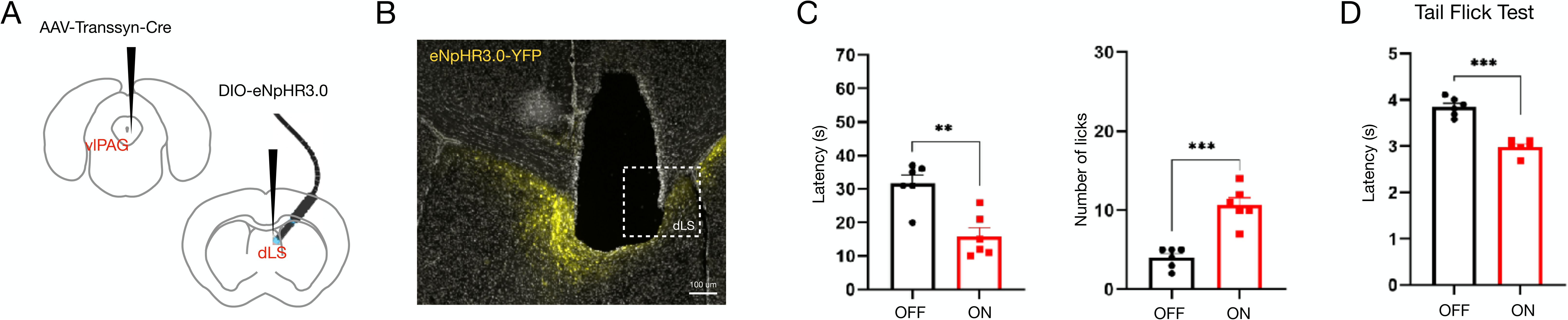
Optogenetic silencing of the dLS_post-vlPAG_ neurons is hyperalgesic. (A) AAVTransyn-Cre was injected in the vlPAG and DIO-eNpHR3.0-YFP in the dLS of wild-type mice with a fiber cannula on the dLS_postvlPAG_ neurons. (B) The expression of eNpHR3.0-YFP (yellow) was confirmed in the dLS_post-vlPAG_ neurons. (C) Latency to lick (seconds) (31.67±2.55 compared to 15.83±2.60, respectively; t-test, \*\**P*=0.0015, n=6) and licks (4.00±0.21 compared to 10.67±4.32, respectively; t-test, \*\*\**P*=0.0001, n=6) on the hot-plate test with yellow light Off and On. (D) Tail-flick latency (seconds) (3.85±0.08 compared to 2.98±0.07, respectively; t-test, \*\*\**P*=0.0001, n=6) with yellow light Off and On.

### dLS_post-vlPAG_ neurons project to LHA

The LS neurons, through their projections to the lateral hypothalamic area (LHA), were shown to mediate RS-induced SIA (Shah et al., 2024). Hence, we wondered if the dLS neurons that innervate the LHA also receive synaptic inputs from the vlPAG. To test that, we performed retrograde monosynaptic rabies tracing (Wickersham et al., 2007; Callaway and Luo, 2015) from the pre-synaptic dLS neurons to LHA (Figure 6A). Retrograde monosynaptic rabies tracing enables unbiased genetic labeling of pre-synaptic neurons. We injected the retrogradely transporting RetroAAV-Cre in the LHA and intersected with Cre-dependent DIO-G and DIO-TVA-GFP in the LS (Figure 6A). After three weeks, we injected the delG-Rabies-mCherry in the dLS (Figure 6A). As expected, we observed the starter cells (yellow) that co-expressed TVA-GFP and delG-Rabies-mCherry in the dLS (Figure 6B). We observed red cell bodies in the vlPAG (Figure 6C) that were retrogradely transported in a transsynaptic manner from the LS neurons pre-synaptic to the LHA. Thus, the neurons in the vlPAG are pre-synaptic to the LS neurons with LHA projections. Next, in a complementary experiment, we injected the AAVTrannsyn-Cre in the vlPAG and the DIO-GFP in the LS to label the dLS_post-vlPAG_ neurons (Figures 6D, E). We observed GFP terminals in the LHA (Figure 6F) as predicted. We simultaneously injected AAV-turboRFP in the LS (Figure 6D, E). This dual labeling strategy labeled the dLS_post-vlPAG_ neurons in green while all LS neurons were in red (Figure 6E). Similarly, the axonal projections of the LS projections to LH and dLS_post-vlPAG_ neurons to LH were labeled green and red, respectively (Figure 6F). The nature of overlap (Figure 6F) indicates that many LS neurons projecting to the LHA receive inputs from the vlPAG. Moreover, this overlap was seen across the range of anterior-posterior coordinates in the mouse LHA (Figure 6G).

**Figure 6.**
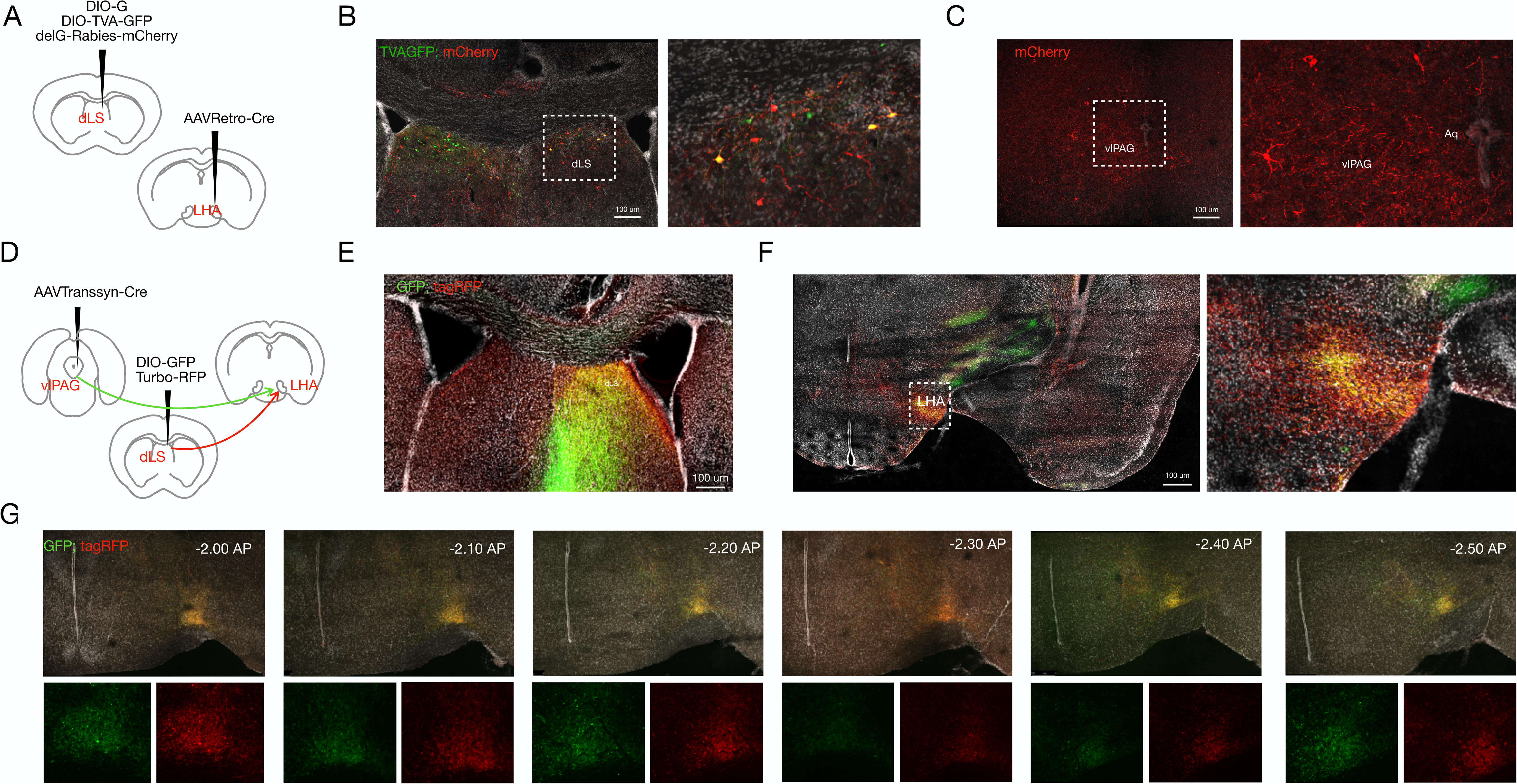
dLS_post-vlPAG_ neurons project to the LHA. (A) AAVRetro-Cre was injected in the LHA, and DIO-G and DIO-TVA-GFP were injected in the dLS of wild-type mice. Three weeks later, the delG-Rabies-mCherry virus was injected into the dLS. (B) Starter cells (yellow) were observed in the dLS (marked region, zoomed-in on the right). (C) mCherry-positive cell bodies (red) are seen in the vlPAG (marked region zoomed in on the right). (D) AAVTranssyn-Cre was injected in the vlPAG, and DIO-GFP and Turbo-RFP were injected in the dLS. (E) An overlap of red and green cells (yellow) is seen in the dLS. (F) An overlap of red and green projections (yellow) is seen in the LHA (marked region zoomed in on the right). (G) Serial rostral to caudal sections of the LHA depicts red and green overlap.

## Discussion

In this study, we have delineated a neural mechanism for how acute restraint, a widely used experimental strategy, induces stress in mice, resulting in short-lasting pain modulation. We found that the vlPAG, a brainstem area involved in escape responses and nociception, provides inputs to the LS (Figures 1B, C, E). The pre-synaptic vlPAG and the post-synaptic LS neurons are tuned to bouts of escape behaviors seen in mice under physical restraint (Figures 2D, 2E, 3C, 3D). Moreover, these neurons respond to noxious heat, and when mice jumped to escape the noxious thermal stimuli, the LS_post-vlPAG_ was engaged (Figures 2G, 2H, 3E, 3F). Notably, bidirectional manipulation of the LS_post-vlPAG_ neurons altered nociceptive thresholds (Figures 4C, 4D, 5C, 5D). Last, the LS_post-vlPAG_ innervated LHA, the downstream target of LS, through which the LS neurons mediate SIA (Figures 6B, 6C, 6E, 6F, 6G).

In the early 1960s, direct electrical stimulation of the PAG was found to be analgesic (Reynolds, 1969). Since then, numerous studies in laboratory animal models across species have established the PAG and the downstream RVM neurons with their inhibitory projections to the dorsal horn to be instrumental in endogenous opioid-induced analgesia (Basbaum and Fields, 1978; Behbehani and Fields, 1979; Shah and Dostrovsky, 1980; Gerhart et al., 1984; Nguyen et al., 2023). Hence, the antinociceptive properties of the PAG are primarily attributed to the downstream connections with the RVM. Importantly, stress-induced analgesia is opioid-dependent and RVM-mediated (Willer et al., 1981; Parikh et al., 2011; Bruehl et al., 2022). Recently, we showed how LS-LHA-RVM circuitry is involved in stress-induced modulation of nociceptive thresholds (Shah et al., 2024). This analgesic circuitry is opioid sensitive and reversible by naloxone. Thus, with the data presented here, we propose that the PAG can modulate nociceptive thresholds and be involved in two concurrent pathways: i) directly through RVM and ii) through LS-LHA-RVM. Physiologically, the PAG-LS pathway may be specific for the RS-induced SIA.

Escape from predators and acutely aversive, including noxious somatosensory stimuli, is critical for survival. The PAG serves as an orchestrator for the escape responses (Branco and Redgrave, 2020). The PAG neurons receive visual and somatosensory stimuli and, when sufficiently aversive, evoke escape from the threatening stimuli. Here, we explored if the escape from acute restraint, which evokes repeated attempts to escape until experimenter-mediated release, is mediated by PAG. Further, we investigated if the PAG neurons’ axonal arborizations into the LS played a role in SIA. SIA is hypothesized to be part of the ensemble of important behaviors for survival. In the context of acute stressors, SIA is an innate mechanism that suppresses pain sensations to deal with the stressors. In the case of RS, it is escaping the restraint. Hence, the vlPAG-LS pathway is engaged in animals undergoing RS, recruiting the downstream neural mechanisms modulating pain. Furthermore, It will be interesting to determine the interoceptive and autonomic signals that activate the PAG neurons in animals under acute restraint.

## Declaration of Competing Interest

The authors declare that they have no known competing financial interests or personal relationships that could have appeared to influence the work reported in this paper.

## List of abbreviations

AP: Anterior-Posterior
BSA: Bovine serum albumin
DCZ: Deschloroclozapine
dLS: dorsal Lateral Septum
DREADD: Designer Receptors Activated Only by Designer Drugs
DV: Dorsal-Ventral
GFP: Green fluorescent protein
i.p.: Intraperitoneal
LHA: Lateral Hypothalamic area
LS: Lateral Septum
ML: Medial-Lateral
NLS: Nuclear localization signal
PAG: Periaqueductal gray
PBS: Phosphate Buffered Saline
PFA: Paraformaldehyde
RFP: Red fluorescent protein
RS: Restraint Stress
SIA: Stress-induced analgesia
SPF: Specific pathogen-free
vlPAG: ventro lateral periaqueductal gray
YFP: Yellow fluorescent protein

## Notes

### Competing Interest Statement

The authors have declared no competing interest.

